# Iterative Refinement of Cellular Identity from Single-Cell Data Using Online Learning

**DOI:** 10.1101/2020.01.16.909861

**Authors:** Chao Gao, Sebastian Preissl, Chongyuan Luo, Rosa Castanon, Justin Sandoval, Angeline Rivkin, Joseph R. Nery, Margarita M. Behrens, Joseph R. Ecker, Bing Ren, Joshua D. Welch

## Abstract

Recent experimental advances have enabled high-throughput single-cell measurement of gene expression, chromatin accessibility and DNA methylation. We previously used integrative non-negative matrix factorization (iNMF) to jointly learn interpretable low-dimensional representations from multiple single-cell datasets using dataset-specific and shared metagene factors. These factors provide a principled, quantitative definition of cellular identity and how it varies across biological contexts. However, datasets exceeding 1 million cells are now widely available, creating computational barriers to scientific discovery. For instance, it is no longer feasible to analyze large datasets using standard pipelines on a personal computer with limited memory capacity. Moreover, there is a need for an algorithm capable of iteratively refining the definition of cellular identity as efforts to create a comprehensive human cell atlas continually sequence new cells.

To address these challenges, we developed an online learning algorithm for integrating large and continually arriving single-cell datasets. We extended previous online learning approaches for NMF to minimize the expected cost of a surrogate function for the iNMF objective. We also derived a novel hierarchical alternating least squares algorithm for iNMF and incorporated it into an efficient online algorithm. Our online approach accesses the training data as mini-batches, decoupling memory usage from dataset size and allowing on-the-fly incorporation of new datasets as they are generated. The online implementation of iNMF converges much more quickly using a fraction of the memory required for the batch implementation, without sacrificing solution quality. Our new approach processes 1.3 million single cells from the entire mouse embryo on a laptop in 25 minutes using less than 500 MB of RAM. We also analyze large datasets without downloading them to disk by streaming them over the internet on demand. Furthermore, we construct a single-cell multi-omic cell atlas of the mouse motor cortex by iteratively incorporating eight single-cell RNA-seq, single-nucleus RNA-seq, single-nucleus ATAC-seq, and single-nucleus DNA methylation datasets generated by the BRAIN Initiative Cell Census Network.

Our approach obviates the need to recompute results each time additional cells are sequenced, dramatically increases convergence speed, and allows processing of datasets too large to fit in memory or on disk. Most importantly, it facilitates continual refinement of cell identity as new single-cell datasets from different biological contexts and data modalities are generated.

## 1 Introduction

### 1.1 Quantitative Definition of Cell Identity from Single-Cell Data

Defining cellular identity is foundational to a genomic approach to medicine, because discovering what goes wrong in disease requires a reference map of the molecular states of healthy cells. Cells have long been qualitatively characterized by a combination of features such as morphology, presence or absence of cell surface proteins, and broad function [1]. Recently, high-throughput single-cell sequencing technologies have enabled researchers to profile multiple molecular modalities, including gene expression, chromatin accessibility and DNA methylation [2]. Integrating multiple single-cell modalities offers tremendous opportunities for unbiased, comprehensive, quantitative definition of discrete cell types and continuous cell states. The resulting catalog of normal cell types promises to revolutionize fields like neuroscience, developmental biology and physiology [3]. Furthermore, knowing the molecular profiles of normal cell types points to biochemical mechanisms by which genetic and environmental factors cause disease.

Multiple features contribute to cell identity, including gene expression, epigenomic modifications, and spatial location within a tissue, but it is not currently possible to simultaneously measure all of these quantities within the same single cells. Experimental methods for assaying transcriptome and epigenome from the same single cells have been demonstrated, but have not been widely adopted due to significant limitations in data quality and/or scalability. Large-scale gene expression, chromatin accessibility, DNA methylation, chromatin conformation, and spatial transcriptomic measurements of different individual cells are now widely available, but these features have generally been used separately to identify cell clusters representing putative cell types, and it is critical to investigate how these different molecular features of cell identity are related [4].

### 1.2 The Need for Scalable Integration of Single-Cell Data

Single-cell data integration thus represents a crucial step toward enabling quantitative definition of cell identity, but existing computational approaches do not address this need. Three unique aspects make single-cell integration challenging: (1) unlike bulk multi-omic data, only one modality is generally available from each single cell; (2) the cell types proportions present in each sample may differ significantly; and (3) number of samples (*n*) per dataset is large and rapidly growing. Currently, scRNA-seq datasets are growing more rapidly than other single-cell data modalities, but we also anticipate rapid growth in the scale of these other data modalities in the near future. Indeed, a recent study assayed more than 100,000 cells with single-cell ATAC-seq [5], and a recent spatial transcriptomic study assayed 1 million cells [6].

A key insight motivating our approach is that techniques for so-called “online learning” [7], in which calculations are performed on-the-fly as new datasets continuously become available (as in many internet applications), provides a path to scalable single-cell data integration.

Several recent single-cell data integration approaches have been developed, including Seurat v3, Harmony, and Scanorama [2, 8, 9], but these approaches are not designed to integrate multiple data types and/or have difficulty scaling to massive datasets. Furthermore, none of these existing methods can incorporate new data, but instead must recalculate results each time new datasets arrive. We address these limitations by developing online iNMF, an algorithm that allows scalable and iterative single-cell multi-omic integration.

### 1.3 Integrative Nonnegative Matrix Factorization

In this paper, we build upon the nonnegative matrix factorization approach at the heart of our recently published LIGER algorithm [10] to develop an online learning algorihm. The intuition behind LIGER is to jointly infer a set of latent factors (“metagenes”) that represent the same biological signals in each dataset, while also retaining the ways in which these signals differ across datasets. These shared and dataset-specific factors can then be used to jointly identify cell types and states, while also identifying and retaining cell-type-specific differences in the metagene features that define cell identities. LIGER starts with two or more single-cell datasets, which may be scRNA-seq experiments across different individuals, time points, or species. The inputs to LIGER may even be measurements from different molecular modalities, such as single-cell epigenome data or spatial transcriptomic data that assay a common set of genes. LIGER relies upon integrative nonnegative matrix factorization (iNMF) [4], which solves the following optimization problem:

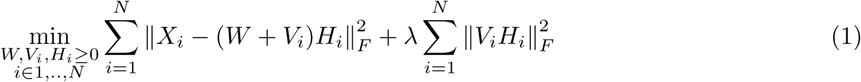

to jointly factorize *N* datasets (each consisting of a genes × cells matrix *X*_*i*_), inferring both shared (*W*) and dataset-specific (*V*_*i*_) factors (**Fig. 1a**). Each factor, or metagene, represents a distinct pattern of gene co-regulation, often corresponding to biologically interpretable signals–like the genes that define a particular cell type. The dataset-specific metagenes (*V*_*i*_) allow robust representation of highly divergent datasets; the factorization can even accommodate missing cell types. Thus, these shared and dataset-specific metagenes can be used to quantitatively define cell identity across biological contexts in terms of inferred co-expressed gene sets or biological pathways.

**Fig. 1.**
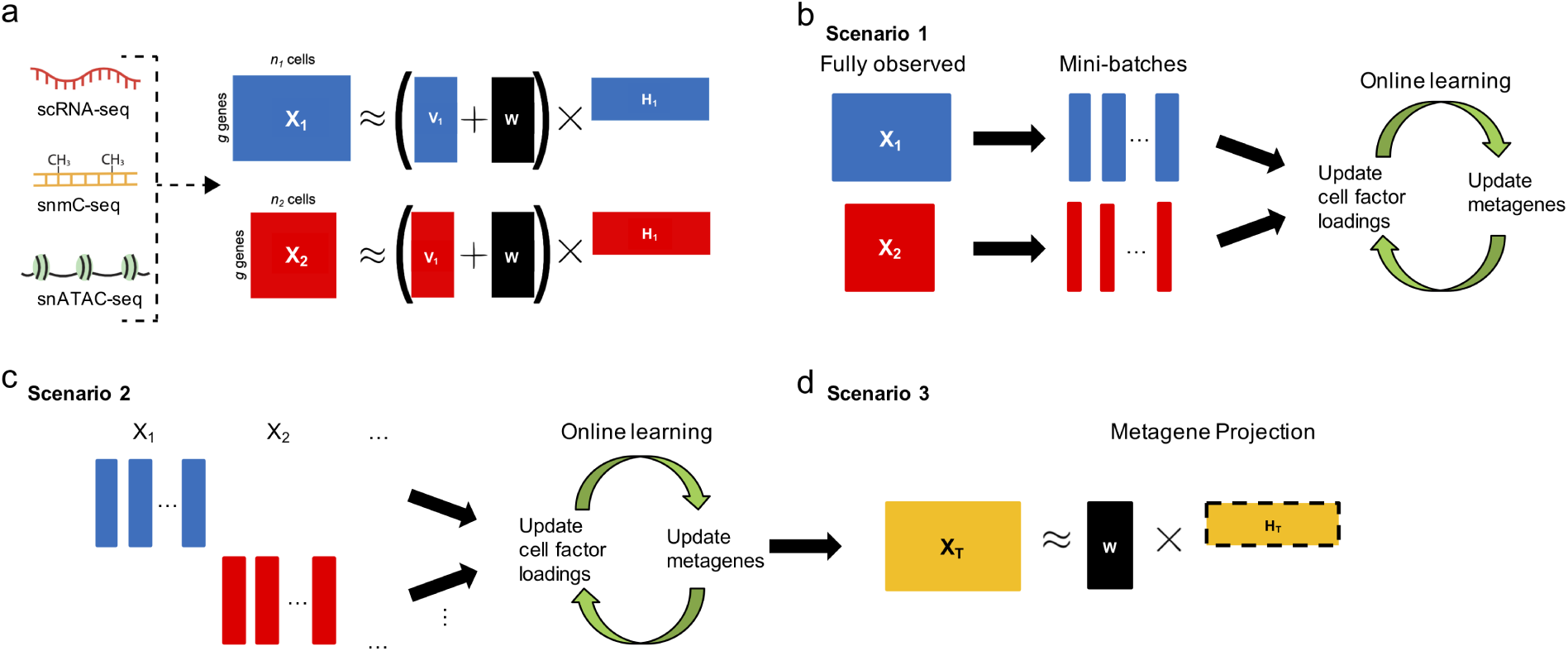
Overview of the online iNMF algorithm. **a**, Schematic of integrative nonnegative matrix factorization (iNMF): the input single-cell datasets are jointly decomposed into shared (*W*) and dataset-specific (*V*_*i*_) metagenes and corresponding “metagene expression levels” or cell factor loadings (*H*_*i*_). These metagenes and their corresponding cell factor loadings provide a quantitative definition of cell identity and how it varies across biological settings. **b-d**, Three different scenarios in which online learning can be used for single-cell data integration. (**b**) Scenario 1: the single-cell datasets are large but fully observed. Online iNMF processes the data in random mini-batches, enabling memory usage and/or disk storage independent of dataset size. Each cell may be used multiple times in different “epochs” of training to update the metagenes. (**c**) Scenario 2: the datasets arrive sequentially, and online iNMF processes the datasets as they arrive, using each cell to update the metagenes exactly once. (**d**) Scenario 3: online iNMF is performed as in scenario 1 or scenario 2 to learn *W* and *V*. Then cell factor loadings for the newly arriving dataset are calculated using the shared metagenes (*W*) learned from previously processed datasets. The new dataset is not used to update the metagenes.

### 1.4 Online Matrix Factorization

Since its proposal by Lee and Seung, nonnegative matrix factorization (NMF) has been widely used to learn interpretable representations of high-dimensional data [11]. NMF is a non-convex optimization problem, so the strongest possible convergence guarantee for an NMF algorithm is that it converges to a local minimum of the objective function. However, the widely used multiplicative update algorithm has no such theoretical convergence guarantee and shows slow convergence in practice. More efficient NMF algorithms based on block coordinate descent, including alternating nonnegative least squares (ANLS) and hierarchical alternating least squares (HALS), have been developed [12], which both converge rapidly in practice and are theoretically guaranteed to converge to a local minimum. Nevertheless, even these approaches are not able to efficiently handle large and streaming inputs such as images and videos (which often arise in web applications, hence the name “online learning”) [13]. As opposed to a batch learning algorithm, an online learning algorithm accesses the data only as either single data points or mini-batches and continually updates the basis elements (metagenes in our context). An online NMF algorithm was developed with guaranteed convergence to a local minimum. This approach showed strong empirical performance and extremely fast convergence compared to batch NMF [7].

### 1.5 Online iNMF

In this study, we extend the online NMF approach of Mairal et al. [7] to make it suitable for iNMF. Online iNMF provides two significant advantages: (1) integration of large multi-modal datasets by cycling through the data multiple times in small mini-batches and (2) integration of continually arriving datasets, where the entire dataset is not available at any point during training (**Fig. 1**).

We envision using online iNMF to integrate single-cell datasets in three different scenarios (**Fig. 1**). In scenario 1, where the datasets are large and fully observed, the algorithm accesses mini-batches from all datasets at the same time and repeatedly updates the metagenes and cell factor loadings. Each cell can be revisited throughout multiple epochs of training (**Fig. 1b**). A key advantage of scenario 1 (compared to batch iNMF) is that only a single mini-batch needs to be in memory at a time. Scenario 1 also allows processing of large datasets without even downloading them on disk, by streaming them over the internet. In scenario 2, the input datasets arrive sequentially, and the online algorithm uses each cell exactly once to update the metagenes, without revisiting data already seen (**Fig. 1c**). The key advantage of scenario 2 is that the factorization is efficiently refined as new data arrives, without requiring expensive recalculation each time. A third scenario allows us to project new data into the latent space already learned, without using the new data to update the metagenes. In scenario 3, we first use online iNMF to learn metagenes as in scenario 1 or scenario 2. Then, we use the shared metagenes (*W*) to calculate cell factor loadings for a new dataset, without using the new data to update the metagenes. This third scenario is highly efficient at incorporating data, allows users to query their data against a curated reference, and provides increased robustness to dataset differences in newly arriving data (**Fig. 1d**).

## 2 Methods

### 2.1 Derivation of Online Algorithm for iNMF

In our previous implementation of iNMF [10], we derived an ANLS algorithm to solve for *H*_*i*_, *W*, and *V*_*i*_. Briefly, ANLS optimizes the iNMF objective by iteratively solving a nonnegative least squares problem to update each of the matrices (*H*_*i*_, *W, V*_*i*_) holding the others fixed. For example, the update for *H*_*i*_ (*i* ∈ {1, …, *N*}) is:

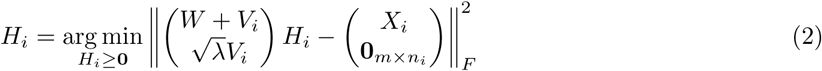

This is a convex nonnegativity-constrained least squares problem that can be solved efficiently using the block principal pivoting algorithm [14]. This ANLS algorithm is guaranteed to converge to a local minimum, and we previously showed that it outperforms the multiplicative updates in practice [10]. We use this strategy for computing the cell factor loadings (*H*) for the cells in the online iNMF algorithm.

Another type of NMF algorithm, hierarchical alternating least squares (HALS), also provides guaranteed convergence to a local minimum, but often shows more efficient convergence in practice [12]. Thus, we sought to derive a novel HALS algorithm for optimizing the iNMF objective. A HALS derivation proceeds by re-writing the objective function as a sum of rank-one approximations (one for each of the *K* inner dimensions of the factorization), then deriving a closed-form solution for each of the *K* basis vectors holding the others fixed. In the case of iNMF, this can be considered a block coordinate descent strategy with (2*N* + 1)*K* vector blocks. For example, for *V*_*i,j*_, the *j*th factor in the dataset-specific metagene matrix *V*_*i*_, we want to solve the optimization problem:

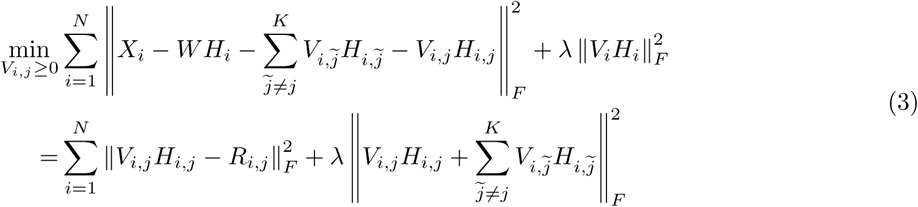

where 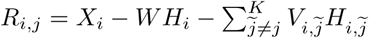.

Taking the derivative with respect to *k*th element in *V*_*i,j*_ gives:

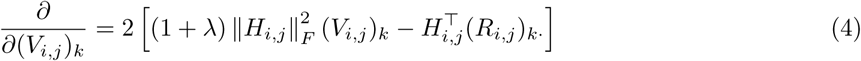

We then set the derivative equal to 0 and solve for (*V*_*i,j*_)_*k*_ subject to nonnegativity constraints. Applying the same process to all elements in *V*_*i,j*_ yields the following update for *j*th column for *V*_*i*_:

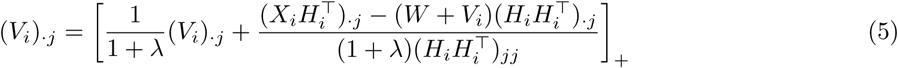

where [*x*]_+_ = max{10^−16^, *x*}. Similar derivation for *W* gives:

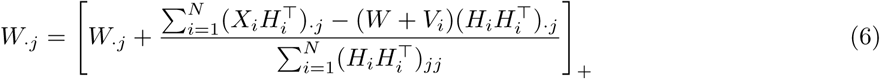

In the online iNMF algorithm, both the shared and dataset-specific metagenes are refined by applying the HALS updates for *W* and *V*_*i*_ to a different mini-batch during each iteration.

### 2.2 Optimizing a Surrogate Function for iNMF

We developed an online learning algorithm for integrative nonnegative matrix factorization by adapting a previously published strategy for online NMF [7]. The key innovation that makes it possible to perform online learning is to optimize a “surrogate function” that asymptotically converges to the same solution as the original iNMF objective. We can formulate NMF using the following objective function: 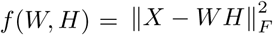 where *W* and *H* are constrained to be nonnegative. The original online NMF paper proved that the following surrogate function converges almost surely to a local minimum as *t* → ∞:

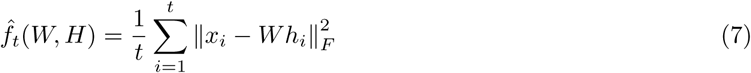

where *H* and *W* are constrained to be nonnegative. We can then perform NMF in an online fashion by iteratively minimizing the expected cost 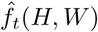 as new data points *x*_*t*_ (or points randomly sampled from a large fixed training set) arrive. Intuitively, this strategy allows online learning because it expresses a formula for incorporating a new observation *x*_*t*_ given the factorization result (*W* ^(*t*−1)^, *H*^(*t*−1)^) for previously seen data points. Thus, we can iterate over the data points one-by-one or in “mini-batches”—and also rapidly update the factorization when new data points arrive. For iNMF, where we have *N* data matrices *X*_1_, …, *X*_*N*_ and data points *x*_*i*_, the iNMF objective function is given by (1). The corresponding surrogate function is:

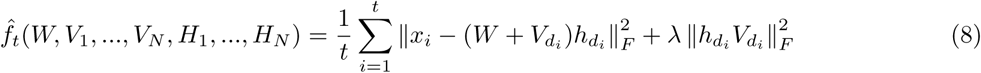

where *d*_*i*_ indicates which dataset the *i*th data point belongs to.

For a new data point (or mini-batch of new data points) *x*_*t*_, we first compute the corresponding cell factor values 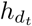. In the original online NMF paper [7], the authors used a least angle regression algorithm (LARS). We chose to use the ANLS update (2) instead because it is highly efficient, designed specifically for NMF (rather than dictionary learning in general) and solves the subproblem exactly in a single iteration. To update the shared (*W*) and dataset-specific (*V*_*i*_) factors, we use the HALS updates (5) and (6), which are analogous to the updates used by Mairal et al [7], but derived specifically for iNMF. Because the updates for *W* and *V*_*i*_ depend on all of the previously seen data points *X*_*i*_ and their cell factor loadings *H*_*i*_, a naive implementation would require storing all of the data and cell factor loadings in memory. However, the HALS updates (5) and (6) depend on *X*_*i*_ and *H*_*i*_ only through the matrix products *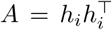* and 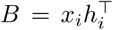. These matrix products have only *K*^2^ and *mK* elements respectively, allowing efficient storage, and can be computed incrementally with the incorporation of each new data point or mini-batch *x*_*t*_:

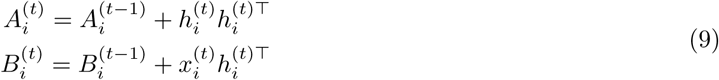

### 2.3 Implementation

We implemented online iNMF according to Algorithm 1 below. We used our previous Rcpp implementation of the block principal pivoting algorithm to calculate the ANLS updates for *h*_*i*_. We implemented the HALS updates for *W* and *V*_*i*_ using native R, since the updates require only matrix operations, which are highly optimized in R. Because the online algorithm does not require all of the data on each iteration (only a single data point or fixed-size mini-batch), we used the `rhdf5` package [15] to load each mini-batch from disk on the fly. By creating HDF5 files with chunk size no larger than the mini-batch size, we were able to create a memory-efficient implementation that never loads more than a single mini-batch of the data from disk at once. In fact, we can even go a step further and analyze datasets that are not stored on the same physical hard drive as the machine performing iNMF. We show below that it is possible to analyze data by streaming over the internet without downloading the entire dataset onto the disk.

For scenario 1, in which the mini-batch size specifies the total number of cells to be processed per iteration across all datasets, we sample *p*_*i*_ cells from each dataset *i*, proportional to its dataset size. Thus, each mini-batch in scenario 1 is composed of a representative sample of cells from all datasets. For scenario 2, in which only one dataset is available at a time, we sample the entire mini-batch from the current dataset. For a mini-batch size of 5,000 cells, reading each mini-batch from disk added minimal overhead (less than 0.35 seconds per iteration) (**Fig. S1**). We also employed two heuristics that were used in the original online NMF paper: (1) we initialized the dataset-specific metagenes using *K* cells randomly sampled from the corresponding input data and (2) we removed information older than two epochs from matrices *A* and *B*.

## 3 Results

### 3.1 Online iNMF Converges Efficiently Without Loss of Accuracy Compared to Batch iNMF

We first assessed the performance of our online iNMF algorithm using two scRNA-seq datasets from the frontal cortex (*n* = 156,167 cells) and posterior cortex (*n* = 99,186 cells) of the adult mouse with 1,111 highly variable genes. These datasets are part of the adult mouse brain atlas recently published by Saunders et al. [16]. Because the online algorithm optimizes the expected cost, we tracked the value of the iNMF objective on both the training data (80% of the entire dataset) and a held-out testing set (20% of the entire dataset) not seen during training. The original online NMF paper also used this evaluation strategy [7]. For this experiment, we set the number of factors (*K*) and the tuning parameter *λ* to 40 and 5, respectively. We compared iNMF against the previously published batch methods for iNMF, including multiplicative updates (Mult) [4] and alternating nonnegative least squares (ANLS) [10]. We calculated *H*_*frontal*_ and *H*_*posterior*_ for the testing set using the *W, V*_*frontal*_ and *V*_*posterior*_ learned on the training set, then used these values to compute the objective on either the training or testing set.

The online iNMF algorithm (mini-batch size = 5,000) converges much faster than ANLS and Multiplicative updates on both the training and held-out set (**Fig. 2a-b**). Within approximately 500 seconds of runtime, the online approach achieves a significantly lower training iNMF objective (**Fig. 2c**). Online iNMF also shows superior performance on several other datasets from different biological contexts (**Fig. S2**). Furthermore, the convergence behavior of the online algorithm on both training and test sets is relatively insensitive to the mini-batch size (**Fig. 2d-e**). For mini-batch sizes from 1,000 to 10,000, the convergence behavior is nearly identical. As the mini-batch size approaches the training dataset size (150,000 or 200,000), the first few iterations take considerably longer, slowing the convergence time, but the final objective remains unchanged. Moreover, for a fixed test set, the runtime needed to reach convergence remains relatively constant once the total number of cells exceeds some minimum threshold (around 50,000, in this case). (**Fig. 2f**). This behavior likely occurs because, for a cell population of fixed complexity (for example, a tissue containing 12 cell types), only some fixed number of observations is required to effectively learn the metagenes. Thus, using the entire dataset to update the shared and data-specific metagenes at each iteration becomes increasingly inefficient as the dataset size exceeds the minimum threshold size needed to learn the metagenes. Conversely, the relative efficiency of online iNMF compared to batch methods increases with dataset size.

**Fig. 2.**
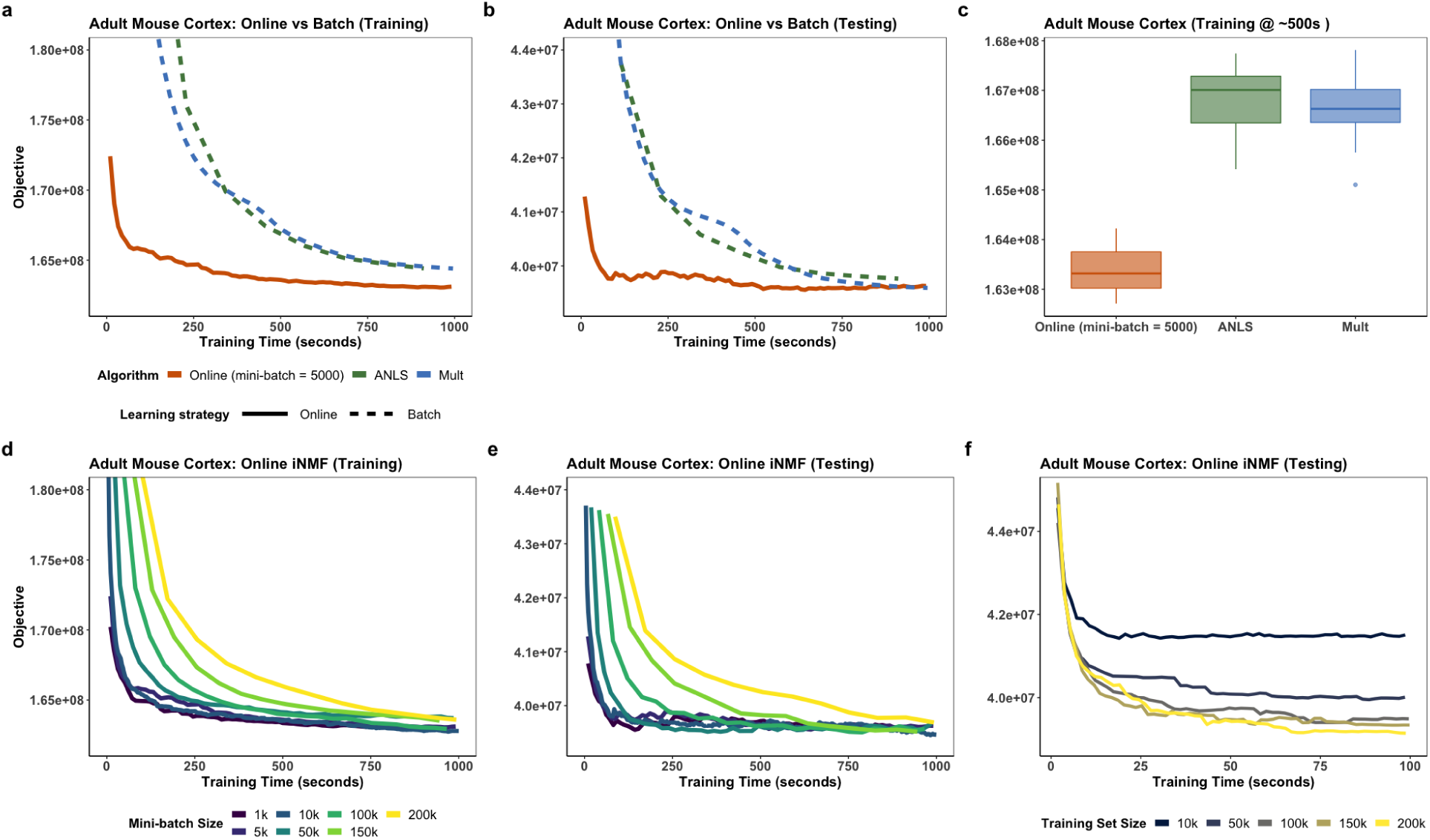
Convergence behavior for online iNMF and two previously published batch iNMF algorithms on scRNA-seq data from the adult mouse cortex. **a,b**, The online iNMF algorithm converges much more rapidly to a similar or better objective function value compared to the previously published batch methods–alternating nonnegative least squares (ANLS) and multiplicative updates (Mult)–on both training and testing sets. **c** The online iNMF algorithm leads to a significantly lower objective function value within the same amount of training time, compared to the batch methods. **d**-**e** The convergence behavior of online iNMF is nearly identical for mini-batch sizes from 1,000 to 10,000. **f**, The online iNMF algorithm becomes increasingly efficient (in terms of decrease in objective function value per unit time) as dataset size increases. The time required for the algorithm to converge does not significantly increase with growing dataset size once the dataset size exceeds 50,000 cells.

#### Algorithm 1 Online Learning for Integrative Nonnegative Matrix Factorization

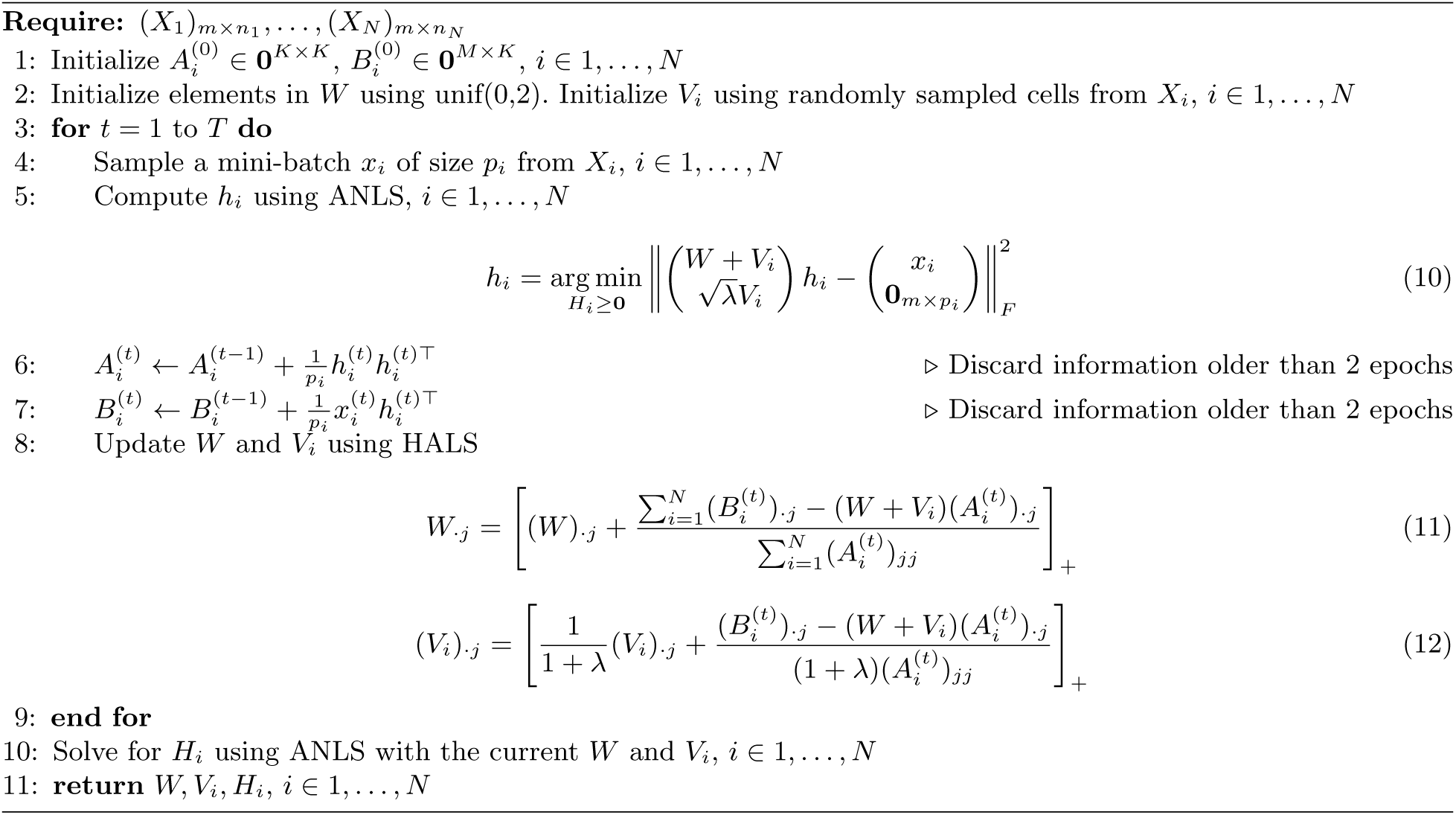

Since ANLS is the batch algorithm that we previously developed for solving the iNMF optimization problem, we refer it as batch iNMF in subsequent analyses. We next investigated whether online iNMF yields similar dataset alignment and cluster preservation to batch iNMF. We applied both online iNMF and batch iNMF to three scRNA-seq data collections, followed by quantile normalization of cell factor loadings. Besides the mouse cortex (*n* = 255,353 cells) datasets mentioned above, the other two datasets are human peripheral blood mononuclear cells (PBMCs, *n* = 13,999 cells) and human pancreatic islets (pancreas, *n* = 14,890 cells). The PBMC dataset consists of 7,451 interferon-*β* (IFNB)-stimulated PBMCs and 6,548 control PBMCs. The human pancreas collection includes eight separate datasets across five sequencing technologies (SMARTSeq2, Fluidigm C1, CelSeq, CelSeq2, and inDrops). For both PBMC and pancreas datasets, we selected 2,000 variable genes for analysis. Then we created UMAP visualizations of the resulting factor loadings, colored by datasets and published cell type labels (**Fig. 3**). These plots allow visual and qualitative assessment of dataset alignment and data structure preservation. As the figure shows, the online iNMF algorithm yields visualizations that are very similar to batch iNMF, suggesting nearly identical dataset alignment and accurate preservation of the original cluster structure for all three data collections.

**Fig. 3.**
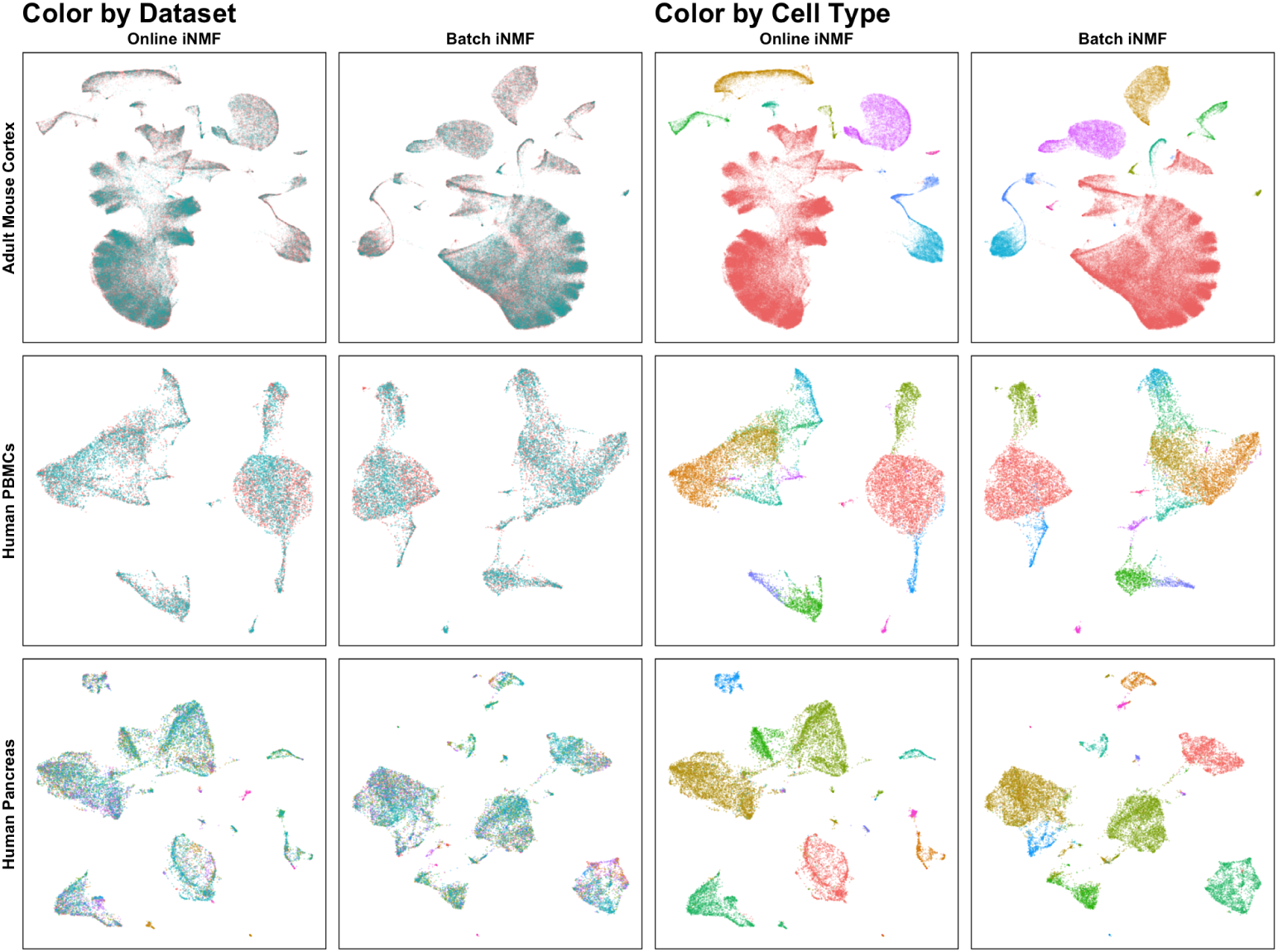
Online and batch iNMF yield highly similar UMAP visualizations. We performed online iNMF and batch iNMF on data from mouse cortex (*n* = 255,353 cells), human PBMCs (*n* = 13,999 cells), and human pancreas (*n* = 14,890 cells). Online iNMF and batch iNMF produce very similar visualizations, suggesting that the approaches give very similar dataset alignment and cluster preservation. We subsequently confirmed this qualitative observation using quantitative metrics (see next section).

### 3.2 Online iNMF Yields State-of-the-Art Single-Cell Data Integration Results Using Significantly Less Time and Memory

We next benchmarked online iNMF (scenario 1) against batch iNMF [10] and two state-of-the-art single-cell data integration methods, Seurat v3 [17] and Harmony [8]. We selected these methods for comparison because a recent paper benchmarked 14 single-cell data integration methods and found that Harmony, Seurat v3 (hereafter referred to as Seurat), and LIGER consistently achieved the best dataset alignment and cluster preservation on a range of datasets [18]. The Harmony algorithm starts from an initial, uncorrected PCA embedding, then integrates scRNA-seq datasets through an iterative process of soft clustering and cluster-specific correction that optimizes for both dataset alignment and cluster separation. The core of the latest Seurat algorithm is canonical correlation analysis (CCA) followed by identification of mutual nearest neighbors (“anchors”). Seurat uses these anchors to calculate batch correction vectors that align the corresponding cells across datasets. Harmony, Seurat, and batch iNMF all require the entire dataset to be stored in memory, so we expect online iNMF to offer substantial improvements in both time and memory usage.

To benchmark time and memory usage, we generated five datasets of increasing sizes (ranging from 10,000 to 255,353 cells in total) sampled from the same adult mouse frontal and posterior cortex data. Then we utilized them to compare the runtime and peak memory usage of online iNMF (mini-batch size = 5,000) and the other methods (**Fig. 4a**).

**Fig. 4.**
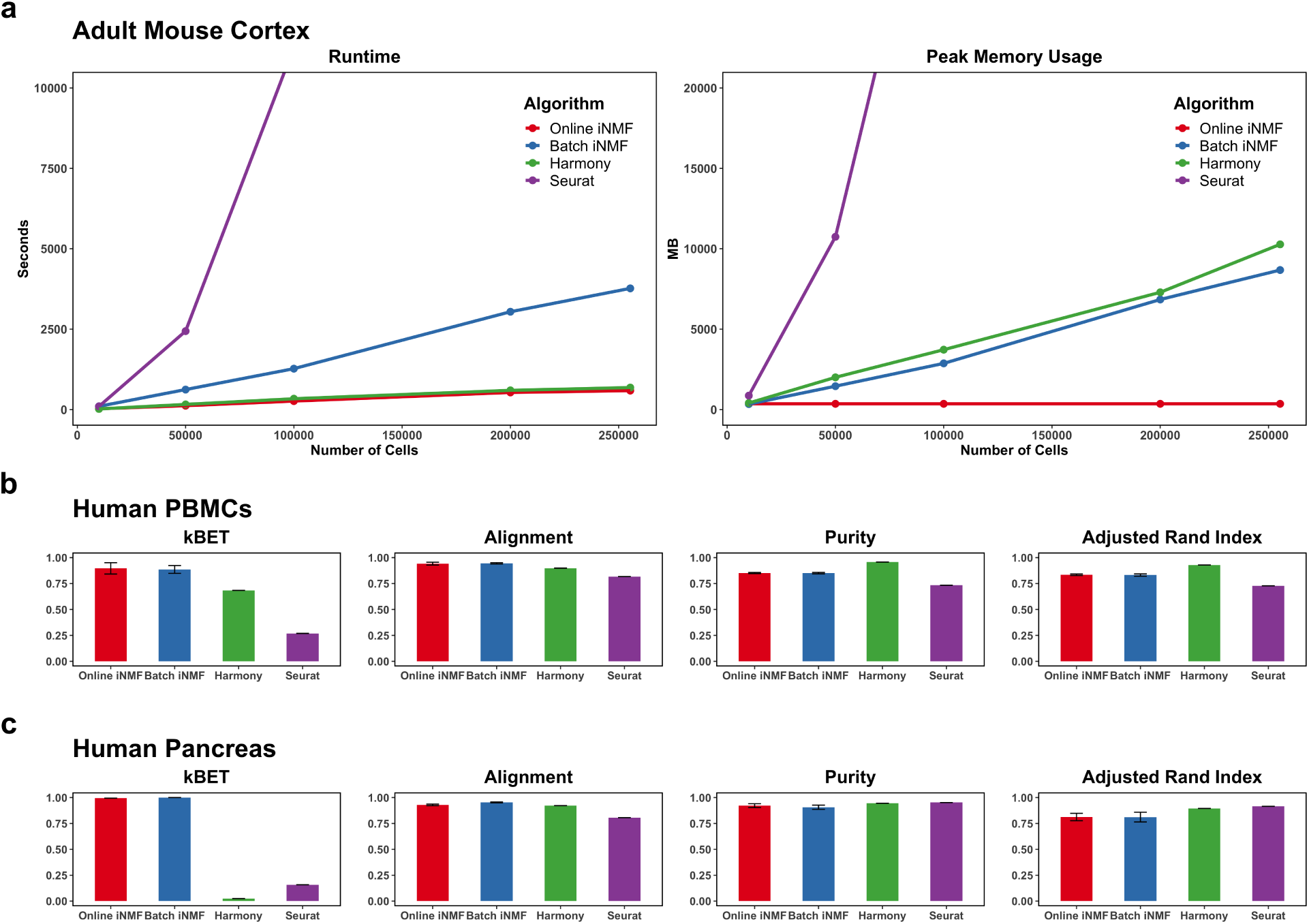
Comparison of convergence time and peak memory usage of online iNMF, batch iNMF, Harmony, and Seurat. The data are sampled from the adult mouse cortex (*n* = 10,000, 50,000, 100,000, 200,000, 255,353 cells, 2 individual datasets), human PBMCs (*n* = 13,999 cells, 2 individual datasets) and human pancreas (*n* = 14,890 cells, 8 individual datasets). **a**, The runtime and peak memory usage required for online iNMF, batch iNMF, Harmony and Seurat to integrate the frontal and posterior cortex datasets. **b, c**, Quantitative assessment of data integration and low-dimensional embedding carried out by four methods on the human PBMCs and human pancreas datasets. Higher values are better for all 4 metrics.

As expected, the runtime required for online iNMF does not increase significantly as the dataset size grows, and the memory usage is constant. Online iNMF is also the fastest method overall, with Harmony the second fastest. Notably, the gap between Harmony and online iNMF widens as the dataset size increases; we also ran both methods on a dataset of 1.3 million cells from the mouse embryo and found that online iNMF finished in 25 minutes using 500 MB of RAM (see below for details), whereas Harmony required 92 minutes and 111 GB of RAM. Seurat and batch iNMF are significantly slower than online iNMF and Harmony on the mouse cortex data, and the runtime of Seurat increases the most rapidly of any method.

Furthermore, the online iNMF algorithm uses far less memory than any other approach, with memory usage completely independent of total data size. By design, online iNMF processes a fixed number of cells during each iteration, which can be determined based on the available computing resources. The peak memory usage of online iNMF with a mini-batch size of 5,000 and *k* = 40 factors is approximately 360MB, no matter how many cells in total are processed. In contrast, batch iNMF, Harmony, and Seurat use increasing amounts of memory as the total number of cells increases. Batch iNMF and Harmony display a similar linear relationship between memory usage and dataset size and use much less memory than Seurat. The memory required by Seurat shows an exceedingly rapid increase given more cells in the input (38 GB of memory is needed for analyzing 100,000 cells). Nevertheless, all three methods besides online iNMF require all of the cells to be loaded in memory, and thus the memory usage grows linearly with dataset size.

Next, we quantified the data integration and cluster preservation performance for online iNMF and the other methods (**Fig. 4b-c**). Following the benchmarking strategy used by Tran et al., we assessed both the alignment performance (measured using two metrics) and the cluster preservation performance (measured using two metrics). For all experiments, we ran both online iNMF and batch iNMF five times, ensuring that the comparisons account for variation due to different initializations. We used the same number of dimensions for all 4 approaches in each comparison (20 dimensions for PBMCs and 40 for pancreas). To quantify dataset alignment, we used the *k*-nearest-neighbor Batch-Effect Test (kBET) metric [19] and the alignment score of Butler et al. [20]. kBET uses a chi-square statistic to test the null hypothesis that the nearest-neighbor distances in the aligned latent space do not differ according to batch. Thus, a higher average p-value indicates better data integration. The alignment score examines the mixing level in the local neighborhood of each cell; proper batch correction produces a large alignment score near 1, while uncorrected datasets have an alignment score near 0. To assess clustering performance, we applied Louvain community detection with the same parameters to the aligned latent spaces obtained by all methods. We then compared the resulting clusters with the published cluster assignments using cluster purity and adjusted Rand index (ARI).

Our results show that online iNMF performs as well as or better than the state-of-the-art methods. The online and batch iNMF algorithms align the PBMCs and pancreas datasets equally well, beating Harmony and Seurat. Furthermore, the online algorithm achieves scores close to batch iNMF on both data collections, confirming the gain in the computational efficiency does not come at the cost of accuracy in data embedding. The difference between iNMF and the other methods is especially pronounced when comparing the values of the kBET metric. We suspect that this difference occurs because our approach includes quantile normalization, which is stronger than the alignment strategies used by Harmony or Seurat. Consistent with our results, the benchmark of Tran et al. also included the pancreas dataset and found that LIGER (batch iNMF) gave substantially higher kBET values than competing methods [18]. The online and batch iNMF algorithms produce comparable clustering results to the other approaches, although Harmony gives slightly higher cluster purity and adjusted rand index. Overall, this analysis indicates that the time and memory efficiency of LIGER does not sacrifice result quality.

We also compared the performance of online iNMF, Seurat, and Harmony when integrating two datasets of different modalities (**Fig. S3**). We used single-nucleus RNA-seq (*n* =101,647 cells) and single-nucleus ATAC-seq (*n* =54,844 cells), which were generated from the mouse primary motor cortex (see next section for more details). Harmony showed the worst alignment performance, possibly because this approach was not originally designed for multi-modal integration, unlike LIGER and Seurat. In contrast, both LIGER and Seurat produced UMAP visualizations indicating successful alignment of RNA and ATAC data. However, the kBET and alignment metrics indicate the LIGER much more thoroughly removes the residual dataset differences than either Seurat or Harmony.

### 3.3 Online iNMF Rapidly Factorizes Large Datasets Using Fixed Memory

To demonstrate the scalability of online iNMF, we applied the algorithm following scenario 1 and analyzed the scRNA-seq data of Saunders et al., which contains 691,962 cells sampled from nine regions (stored in 9 individual datasets) spanning the entire mouse brain. We identified 2,384 genes that are highly variable in at least one of the regions. Using these genes, we performed 3 epochs of iNMF with mini-batch size of 5,000, *K* = 40 and *λ* = 5. We found that quantile normalization was not necessary for this dataset–iNMF alone was sufficient to integrate the datasets. Using online iNMF, we factorized the entire dataset in 24 minutes on a MacBook Pro (Intel i7 processor) using about 350 MB of RAM. We note that the published analysis by Saunders et al. did not analyze all nine tissues simultaneously due to computational limitations. Furthermore, we estimate (based on the data in **Fig. 4a**) that performing this analysis using our previous batch iNMF (ANLS) algorithm would have taken 3 hours and required over 20 GB of RAM.

We then visualized the embedded cells, colored by published cell type labels, in the first and second UMAP coordinates separately for all nine mouse brain regions (**Fig. 5a**). The online iNMF algorithm clearly retains the data structure as the cells within each class are well grouped together. For cells broadly marked as neurons, the distribution varies across regions, indicating neuronal subtypes specialized to different parts of the brain. For example, neurogenic cell populations are identified predominantly in the hippocampus and striatum, consistent with reports of hippocampal and striatal neurogenesis in adult mammals [16, 21, 22].

**Fig. 5.**
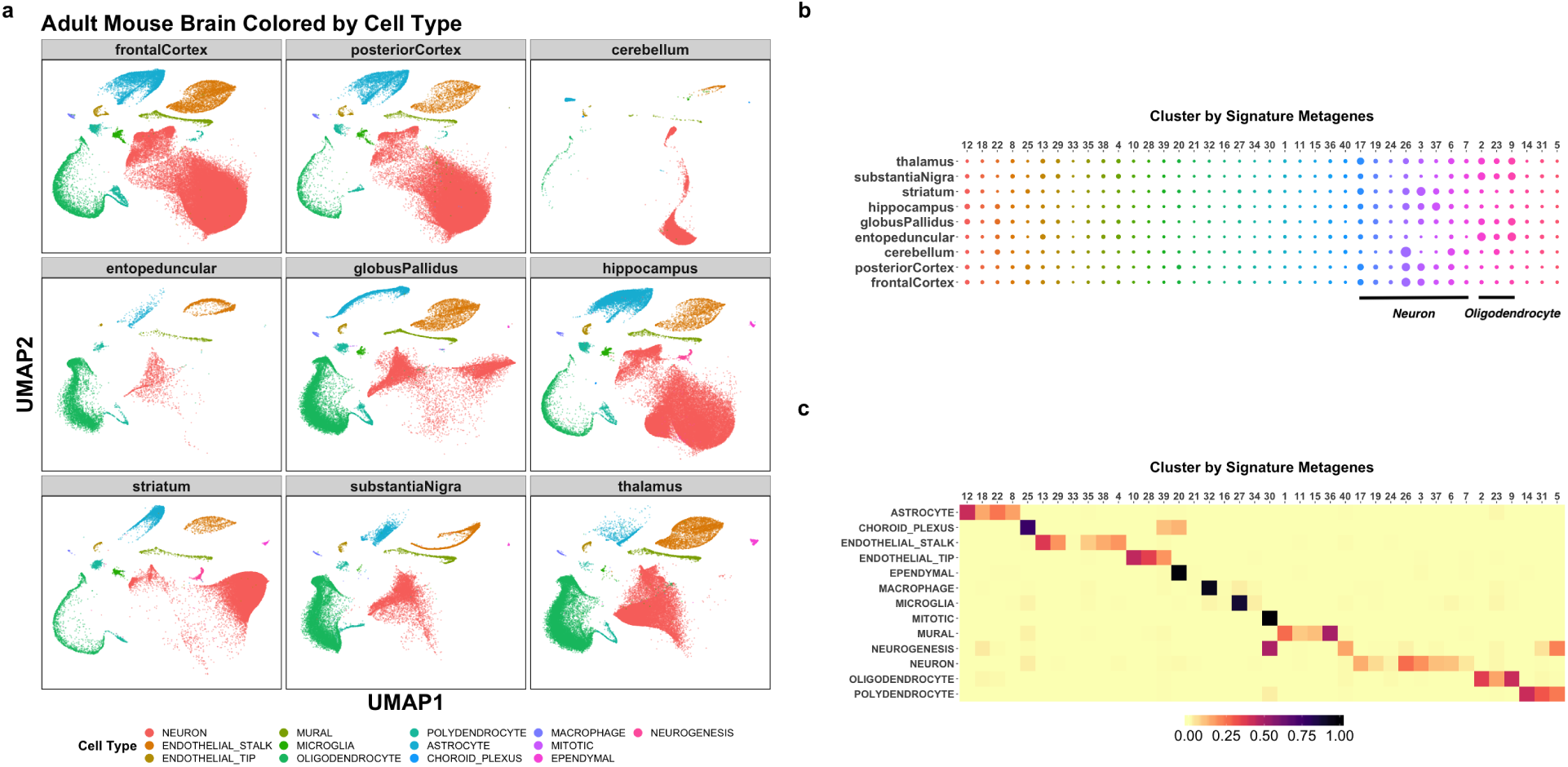
Joint analysis of nine regions of the adult mouse brain (n = 691,962 cells) using online iNMF. **a**, UMAP visualization of the iNMF factors learned for each brain region, colored by published cell class. **b**, Dot plot showing the proportion of each of 40 clusters inferred from iNMF in each brain region. **c**, The heatmap shows the proportion of cells from each cluster in every cell type. The cells in each cluster mostly correspond to a single cell type.

Additionally, we used the factorization to group the cells into 40 clusters by assigning each cell to the factor on which it has the largest loading. We then examined differences in the regional proportions of each cell cluster. Neurons and oligodendrocytes show the most regional variation in composition, consistent with previous analyses [23]. The total proportion of oligodendrocytes varies by region, but individual subtypes of oligodendrocytes are not region-specific, as expected. In contrast, individual subtypes of neurons are highly region-specific, reflecting diverse regional specializations in neuronal function (**Fig. 5b**). We also investigated the biological properties of our cell factor loadings. Reassuringly, our cluster assignments largely represent subtypes within the broad cell classes and do not span class boundaries. As expected, neurons show by far the most diversity with eight subclusters. In contrast, ependymal cells, macrophages, microglia, and mitotic cells each correspond to only a single cluster (**Fig. 5c**).

To further demonstrate the scalability of online iNMF, we analyzed the mouse organogenesis cell atlas (MOCA) recently published by Cao et al. [24]. After filtering, MOCA consists of transcriptomic profiles for 1,363,063 cells from embryos between 9.5 to 13.5 days of gestation. Using 2,557 highly variable genes, we performed online iNMF in 25 min using less than 500MB of RAM (mini-batch size = 5,000, *K* = 50, *λ* = 5, epoch = 1). Note that the memory usage is higher for MOCA than for the mouse brain dataset because of the higher value of *k*, not because of the number of cells. We then performed a 3D UMAP embedding using the same parameters chosen by Cao et al. [24]. The resulting visualization shows that the cells from all five gestational ages are well aligned (**Fig. S3a**), and the structure of 10 different developmental trajectories as defined by Cao et al. is also accurately preserved (**Fig. S3b**).

We reasoned that, because online iNMF processes only one mini-batch at a time, our approach allows processing datasets by streaming them over the internet, removing the need to store multiple copies of large datasets. To demonstrate this capability, we created an HDF5 file containing the mouse cortex datasets (*n* = 255,353 cells) and saved the file on a remote server maintained by the HDF group. Using the remote HDF5 capabilities of the `h5pyd` package, we read mini-batches over the internet, rather than from disk. Processing the cortex dataset in this fashion took about 18 minutes for completing iNMF, compared to around 6 minutes using local disk reads. This capability provides the unique advantage that users can re-analyze large cell atlases, without requiring each user to download and store the entire data collection.

### 3.4 Online allows Iterative Refinement of Cell Atlas from Mouse Motor Cortex

One of the most appealing properties of our online learning algorithm is the ability to incorporate new data points as they arrive. This capability is especially useful for large, distributed collaborative efforts to construct comprehensive cell atlases [25, 26, 3]. Such cell atlas projects involve multiple research groups asynchronously generating experimental data with constantly evolving protocols, making the ultimate cell type definition a moving target.

To demonstrate the utility of online iNMF for iteratively refining cell type definitions, we used data generated by the BRAIN Initiative Cell Census Network (BICCN), an NIH-funded consortium that aims to identify all of the cell types in the mouse and human brains. During a pilot phase starting in 2018, the BICCN generated single-cell datasets from a single region of mouse brain (primary motor cortex, MOp) spanning 4 modalities (single-cell RNA-seq, single-nucleus RNA-seq, single-nucleus ATAC-seq, single-nucleus methylcytosine-seq) and totaling 786,605 cells. These datasets have been publicly released on the BICCN data portal (https://nemoarchive.org/). Over the past two years, the four consortium centers have sequentially generated datasets, re-running the experiments as additional replicates and new protocols become available. Thus, this data collection provides an ideal case study to demonstrate how online iNMF can refine a cell atlas as additional cells are sequenced.

Following scenario 2 (**Fig. 1c**), we used online iNMF to incorporate the MOp datasets (*n* = 408,885 neurons from eight datasets after filtering, 3717 selected genes) in chronological order, refining the factorization with each additional dataset (**Fig. 6**). These datasets represent a sort of historical record reflecting the rapid development of single-cell experimental techniques, with the first dataset generated using SMART-seq, the dominant protocol before the advent of droplet-based protocols. Subsequent datasets reflect newer technologies, including two versions of the 10X Genomics scRNA-seq protocol (v2 and v3); droplet-based single-nucleus RNA-seq; droplet-based single-nucleus ATAC-seq; and single-nucleus methylcytosine-seq. We used a fixed mini-batch size of 5,000 cells, *k* = 40, *λ* = 5, and performed a single epoch of training (each cell participates in exactly one mini-batch). When adding a new dataset *i*(*i* ≥ 1), we incorporated a new dataset-specific metagene *V*_*i*_ and randomly initialized it. We did not use the previously seen data to refine the factors after the initial single epoch per dataset. Then we re-computed the cell factor loadings for all datasets (*H*_1_, …, *H*_*i*_) using the latest metagenes. Lastly, we quantile normalized these cell factor loadings.

**Fig. 6.**
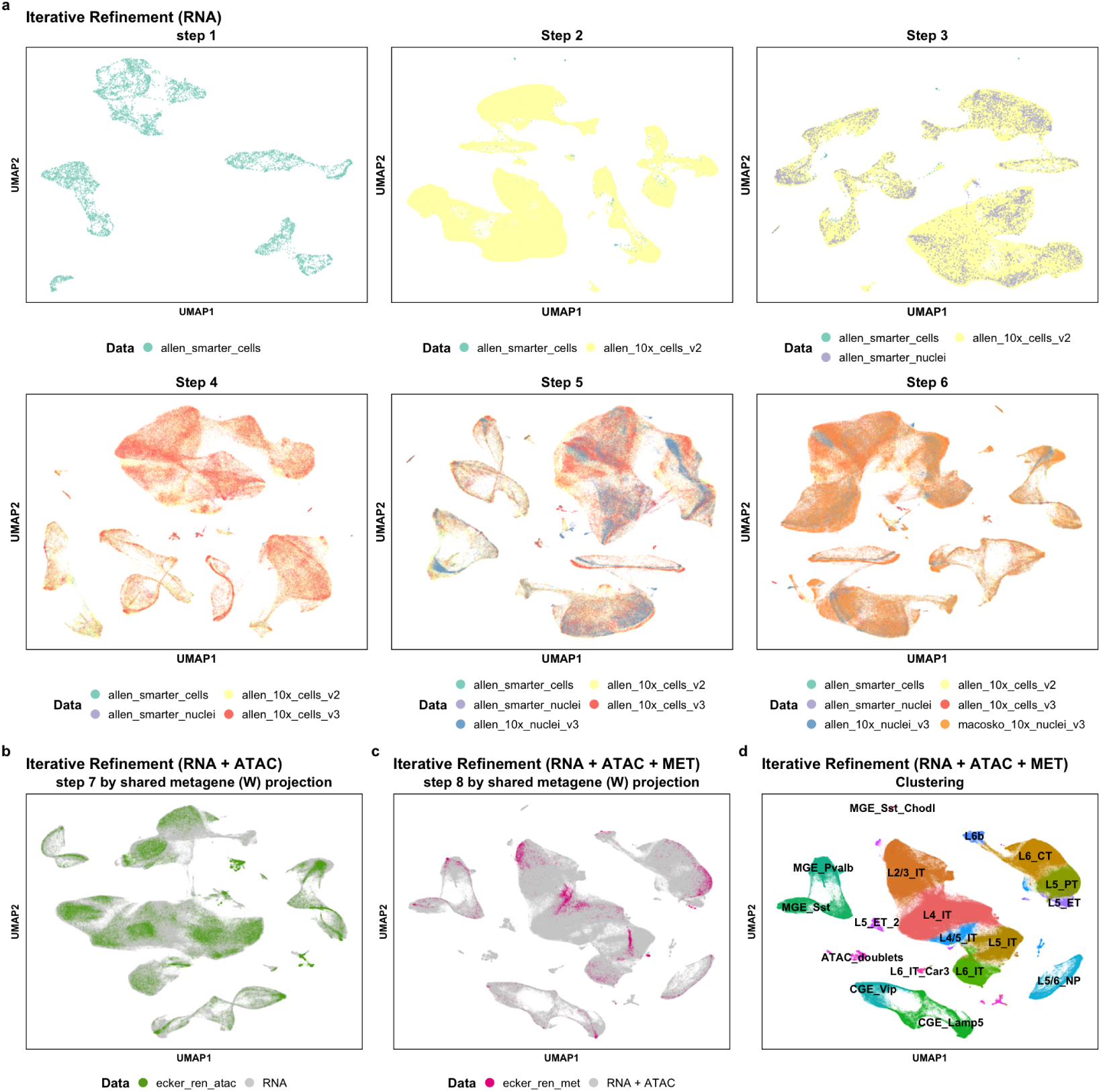
Iterative refinement of cell identity using multiple single-cell modalities from mouse primary motor cortex. We integrated four scRNA-seq datasets, two snRNA-seq datasets, one snATAC-seq dataset and one snmC-seq dataset (*n* = 408,885 neurons). **a**, Sequential integration of six scRNA-seq datasets (scenario 2). **b**, Integration of snATAC-seq data in addition to the scRNA-seq and snRNA-seq data using the shared metagenes (*W*) learned in **(a)** (scenario 3). **c**, Integration of DNA methylation data (snmC-seq) by shared metagene projection (scenario 3). **d**, Joint clusters obtained using the cell factor loadings of all eight aligned datasets. The clusters were annotated based on the visual cortex cell types from Tasic et al.

Our approach successfully incorporates each new single-cell or single-nucleus RNA-seq dataset without revisiting previously processed cells (**Fig. 6a**). Although no ground truth labels or published clustering assignments are available for this data collection, UMAP visualizations indicate that the structure of the datasets is iteratively refined with each successive dataset that is added. However, the single-nucleus ATAC-seq dataset, the 7th dataset to arrive, does not align as well as the RNA datasets when processed according to scenario 2 (**Fig. S5**). We reasoned that this may be because the ATAC-seq data is a completely different modality, and thus scenario 2 may not be the best strategy for incorporating the ATAC-seq. Thus, we also integrated the scATAC-seq data according to scenario 3 (**Fig. 1**): We first performed online iNMF on the MOp scRNA-seq data, then used the shared metagenes (*W*) to project the snATAC-seq data into the same latent space as the scRNA-seq and snRNA-seq data (**Fig. 6b**). Then we integrated the snmC-seq data in the same way (**Fig. 6c**). This strategy produced excellent integration results, and we were able to jointly identify 17 cell types from the scRNA-seq, snRNA-seq, snATAC-seq and snmC-seq data (**Fig. 6d**). Using marker gene expression, we labeled these cell types according to the taxonomy for mouse visual cortex neurons recently published by Tasic et al [27].

We also confirmed that scenario 2 is robust to the order of dataset arrival. To do this, we inspected the effect of random initializations and random ordering of the input datasets on the iterative refinement of cell identity (scenario 2). We ran the online algorithm on the six RNA datasets with five different initializations and five different dataset orders. Different orders result in comparable variation in final cluster assignments compared to the variation from random initialization (average pairwise adjusted Rand index = 0.706±0.049 from random input orders vs. 0.693±0.071 from random initializations). Additionally, UMAP visualizations colored by our final cell type annotations are qualitatively very similar.(**Fig. S6**).

## 4 Discussion

The online iNMF algorithm processes single-cell datasets, possibly from different modalities, each assaying a common set of genes. By reading mini-batches from disk, online iNMF not only converges faster than batch approaches, but also decouples memory usage from dataset size. The online algorithm can even process large datasets stored on a remote server without the need to keep the datasets on a local disk. We anticipate that the efficiency gains of online iNMF will be even greater as the scale of single-cell datasets increases. Furthermore, we do not sacrifice performance for efficiency–our online algorithm performs as well as or better than state-of-the-art methods, including batch iNMF, Harmony and Seurat.

We envision online iNMF enabling single-cell data integration in three different scenarios. In scenario 1, when all single-cell datasets are currently available, the online iNMF algorithm rapidly factorizes the single-cell data into metagenes and cell factor loadings using multiple epochs of training, as we demonstrated on the adult mouse brain data (nine regions) collection. In scenario 2, the online algorithm iteratively refines cell identity as the single-cell datasets sequentially arrive. We demonstrated this capability using single-cell gene expression data from mouse primary motor cortex. We anticipate that scenario 2 will prove useful as researchers continually incorporate newly sequenced cells to build comprehensive cell atlases. Scenario 3 proved especially useful for integrating completely different data modalities, such as the single-cell RNA-seq, single-nucleus ATAC-seq, and single-nucleus DNA methylation data from the mouse primary motor cortex.

As more single-cell datasets of rapidly increasing size become available, online iNMF holds great promise for integrating single-cell multi-omic datasets and cataloging cell identity using limited computational resources.

## Acknowledgments

This work was supported by NIH grants R01 AI149669-01 R01 HG010883-01 (JDW) and 5U19MH114831 (JRE); JRE is an Investigator of the Howard Hughes Medical Institute.

## Author Contributions

SP, CL, RC, JS, AR, JRN, MMB, JRE, and BR generated the snATAC-seq and snmC-seq data. JDW conceived the idea of online iNMF. CG and JDW developed the online iNMF algorithm. CG and JDW wrote the paper. All authors read and approved the final manuscript.

## Software Availability

An R package implementing the LIGER algorithm is available at https://github.com/MacoskoLab/liger

## Supplementary

**Fig. S1.**
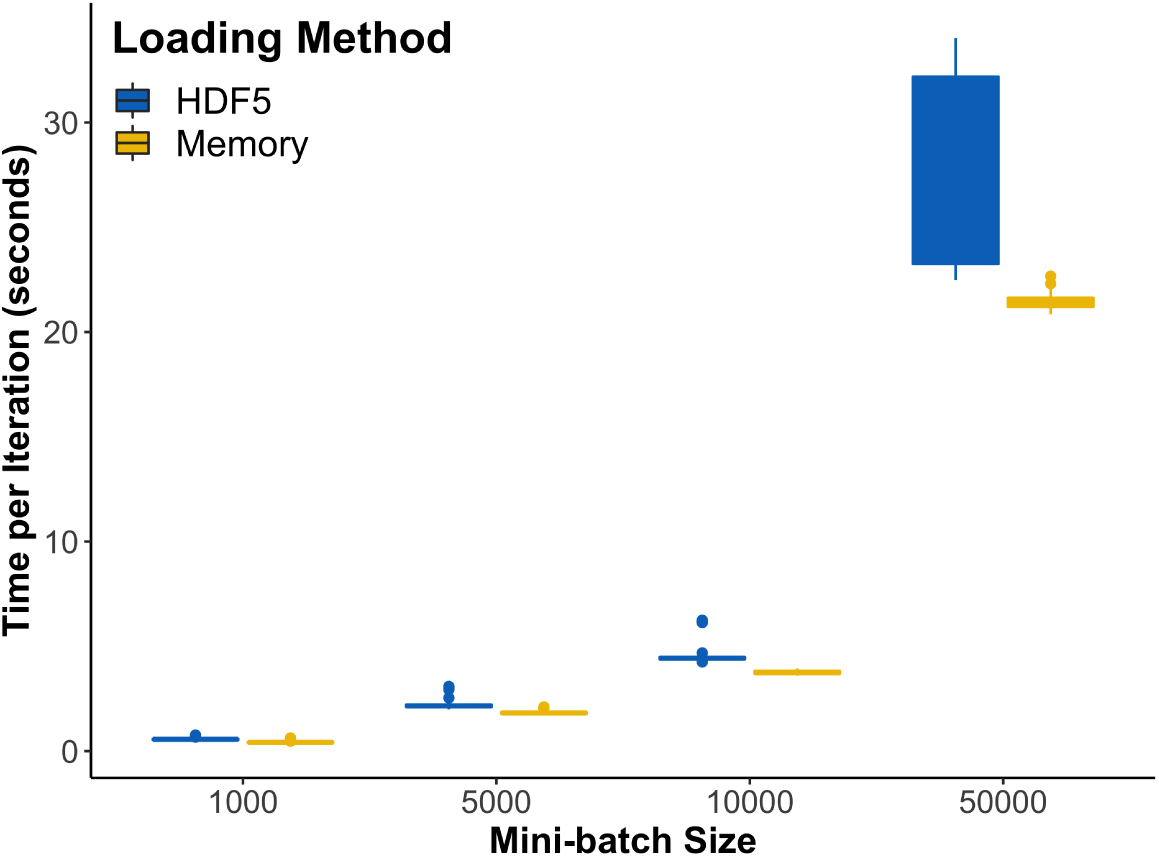
Reading mini-batches from disk adds minimal overhead. In this study, each chunk in HDF5 files stores 1000 samples. Pulling data from the disk does not add significant overhead compared to loading the data from memory, as long as the mini-batch size is close to the specified chunk size.

**Fig. S2.**
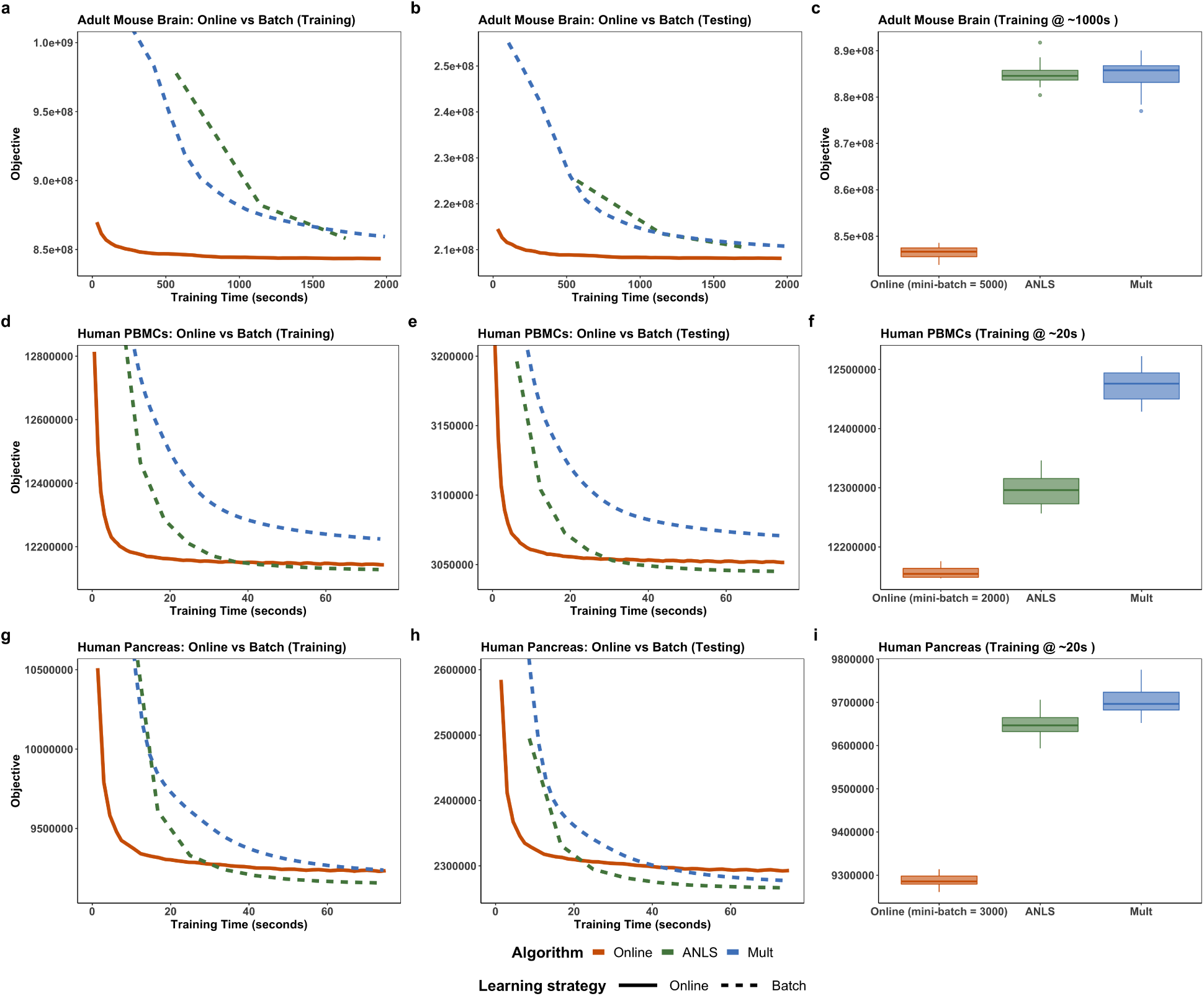
Convergence behavior for online iNMF and two batch iNMF algorithms on scRNA-seq data from the adult mouse brain, human PBMCs and human pancreas. The online iNMF algorithm exhibits faster convergence and better objective minimization shortly after a fixed amount of training time. The advantage of the online algorithm in convergence speed is more apparent for larger datasets. **a**-**c** Adult mouse brain (*n* = 691,962 cells, 9 individual datasets). **d**-**f** Human PBMCS (*n* = 13,999 cells, 2 individual datasets). **g**-**i** Human pancreas (*n* = 14,890 cells, 8 individual datasets).

**Fig. S3.**
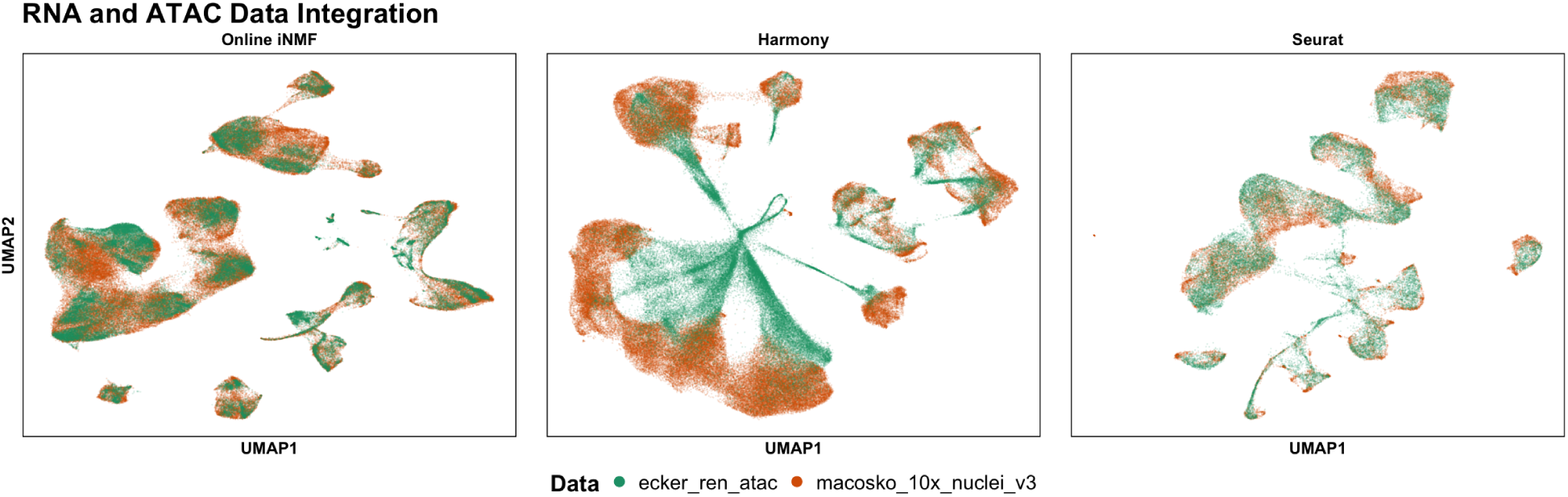
Benchmarking integration using datasets of different modalities. Online iNMF efficiently and accurately integrates scRNA-seq (*n* = 101,647 cells) and scATAC-seq (*n* = 54,844 cells) datasets from the mouse primary motor cortex (kBET = 0.923, Alignment score = 0.737). Harmony does not align the two modalities as well as either Seurat or online iNMF (kBET = 0, Alignment score = 0.305). Seurat aligns the datasets, but shows some residual dataset differences (kBET = 0, Alignment score = 0.645) (*n* = 25,000 cells from each dataset due to memory constraints).

**Fig. S4.**
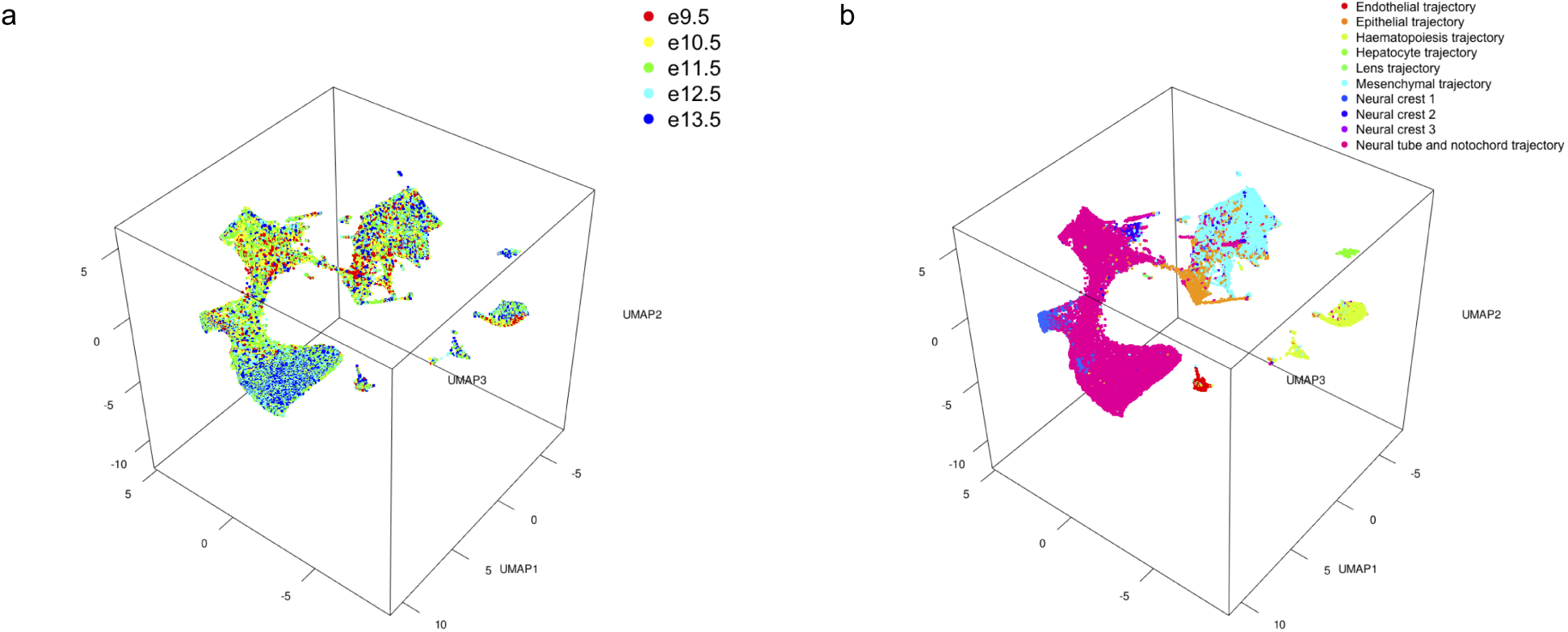
Online iNMF efficiently factorizes the mouse organogenesis cell atlas (MOCA). The MOCA dataset consists of *n* = 1,363,063 cells from embryos between 9.5 to 13.5 days of gestation. The online iNMF analysis required 25 minutes and less than 500 MB of RAM on a MacBook Pro, compared to 92 minutes and 111 GB of RAM for Harmony. **a-b**, 3D UMAP visualization of the online iNMF results, colored by dataset **(a)** and published developmental trajectory labels **(b)**.

**Fig. S5.**
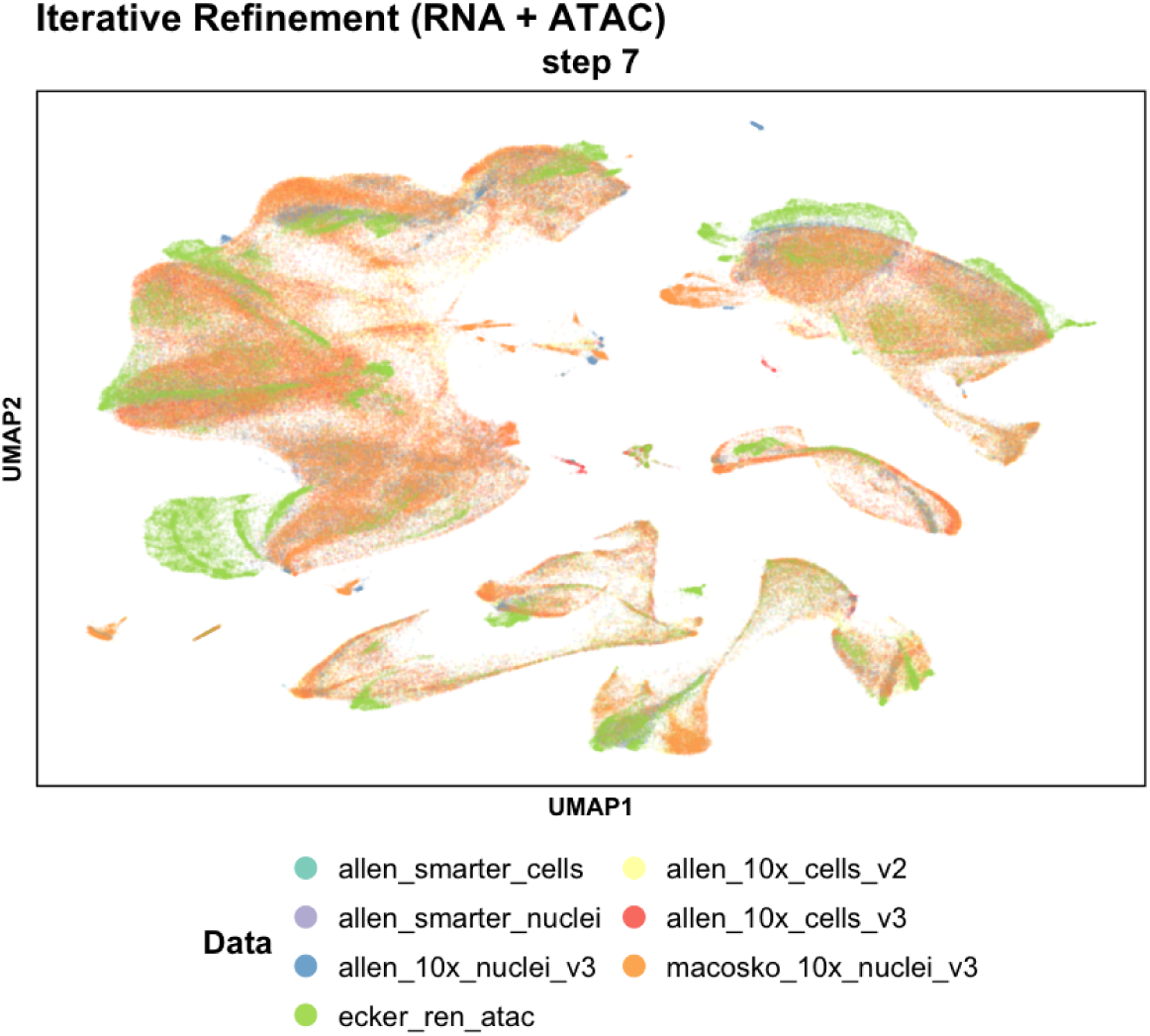
Integration of single-cell RNA-seq and ATAC-seq datasets in scenario 2. Six RNA (scRNA-seq, snRNA-seq) datasets and one ATAC-seq dataset from the mouse primary motor cortex are aligned through iterative refinement.

**Fig. S6.**
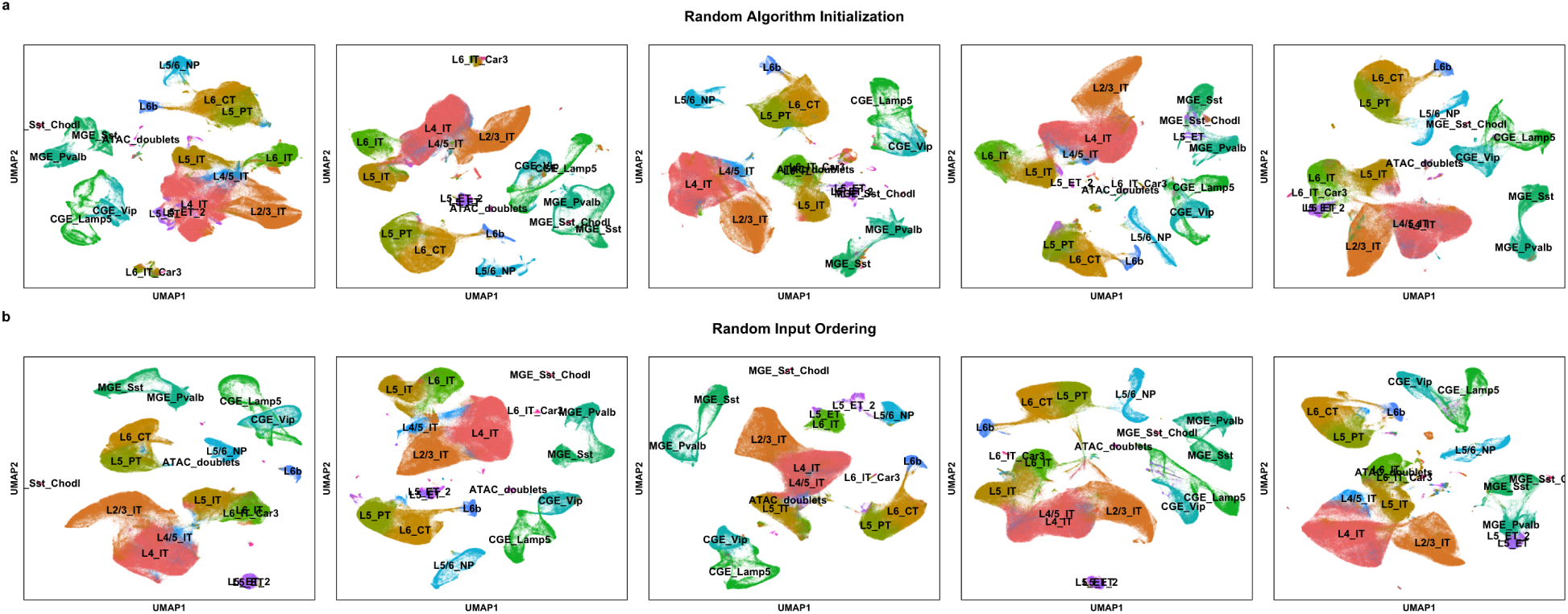
Effect of random algorithm initialization and random input ordering on iterative refinement of cell identity. Online iNMF are applied five times on six MOp scRNA-seq and snRNA-seq datasets for each random setting (scenario 3). **a**, random algorithm initialization. **b**, random input ordering. Clusters from random orderings had an average pairwise ARI of 0.706 (±0.049), compared to an average pairwise ARI of 0.693 (±0.071) for random initializations.

## Notes

#### Summary of Updates

Data generating co-authors added. Benchmarking analyses and additional datasets incorporated.

